# Increased connectivity among sensory and motor regions during visual and audiovisual speech perception

**DOI:** 10.1101/2020.12.15.422726

**Authors:** Jonathan E. Peelle, Brent Spehar, Michael S. Jones, Sarah McConkey, Joel Myerson, Sandra Hale, Mitchell S. Sommers, Nancy Tye-Murray

## Abstract

In everyday conversation, we usually process the talker’s face as well as the sound of their voice. Access to visual speech information is particularly useful when the auditory signal is degraded. Here we used fMRI to monitor brain activity while adult humans (n = 60) were presented with visual-only, auditory-only, and audiovisual words. The audiovisual words were presented in quiet and several signal-to-noise ratios. As expected, audiovisual speech perception recruited both auditory and visual cortex, with some evidence for increased recruitment of premotor cortex in some conditions (including in substantial background noise). We then investigated neural connectivity using psychophysiological interaction (PPI) analysis with seed regions in both primary auditory cortex and primary visual cortex. Connectivity between auditory and visual cortices was stronger in audiovisual conditions than in unimodal conditions, including a wide network of regions in posterior temporal cortex and prefrontal cortex. In addition to whole-brain analyses, we also conducted a region-of-interest analysis on the left posterior superior temporal sulcus (pSTS), implicated in many previous studies of audiovisual speech perception. We found evidence for both activity and effective connectivity in pSTS for visual-only and audiovisual speech, although these were not significant in whole-brain analyses. Taken together, our results suggest a prominent role for cross-region synchronization in understanding both visual-only and audiovisual speech that complements activity in “integrative” brain regions like pSTS.

## Introduction

Understanding speech in the presence of background noise is notoriously challenging, and when visual speech information is available, listeners make use of it—performance on audiovisual (AV) speech in noise is better than for auditory-only speech in noise (Sumby and Pollack, 1954). Although there is consensus that listeners make use of visual information during speech perception, there is little agreement either on the neural mechanisms that support visual speech processing or on the way in which visual and auditory speech information are combined during audiovisual speech perception.

One longstanding perspective on audiovisual speech has been that auditory and visual information are processed through separate channels, and then integrated at a separate processing stage (Grant and Seitz, 1998; Massaro and Palmer, 1998). Audiovisual integration is thus often considered an individual ability that some people are better at and some people are worse at, regardless of their unimodal processing abilities (Magnotti and Beauchamp, 2015; Mallick et al., 2015).

However, more recent data have brought this traditional view into question. For example, Tye-Murray and colleagues (2016) showed that unimodal auditory-only and visual-only word recognition scores accurately predicted AV performance, and factor analyses revealed two unimodal ability factors with no evidence of a separate integrative ability factor. These findings suggest that rather than a separate stage of audiovisual integration, AV speech perception may depend most strongly on the coordination of auditory and visual inputs (Sommers, 2021).

Theoretical perspectives on audiovisual integration have also informed cognitive neuroscience approaches to AV speech perception. Prior functional neuroimaging studies of audiovisual speech processing have largely focused on identifying brain regions supporting integration. One possibility is that the posterior superior temporal sulcus (pSTS) combines auditory and visual information during speech perception. The pSTS is anatomically positioned between auditory cortex and visual cortex, and has the functional properties of a multisensory convergence zone (Beauchamp et al., 2004). During many audiovisual tasks, the pSTS is differentially activated by matching and mis-matching auditory-visual information, consistent with a role in integration (Stevenson and James, 2009). Moreover, functional connectivity between the pSTS and primary sensory regions varies with the reliability of the information in a modality (Nath and Beauchamp, 2011), suggesting that the role of the pSTS may be related to combining or weighing information from different senses.

A complementary proposal is that regions of premotor cortex responsible for representing articulatory information are engaged in processing speech (Okada and Hickok, 2009). The contribution of motor regions to speech perception is hotly debated. Evidence consistent with a motor contribution includes a self-advantage in both visual-only and AV speech perception (Tye-Murray et al., 2013, 2015), and effects of visual speech training on speech production (Fridriksson et al., 2009; Venezia et al., 2016). However, premotor activity is not consistently observed in neuroimaging studies of speech perception, and in some instances may also reflect non-perceptual processing (Szenkovits et al., 2012; Nuttall et al., 2016). It is also possible that premotor regions are only engaged in certain types of speech perception situations (for example, when there is substantial background noise, or when lipreading); individual differences in hearing sensitivity or lipreading ability also may affect the involvement of premotor cortex.

In addition to looking for brain regions that support visual-only or AV speech perception, we therefore broaden our approach to study the role played by effective connectivity between auditory, visual, and motor regions. If a dedicated brain region is necessary to combine auditory and visual speech information, we would expect to see it active during audiovisual speech. If changes in effective connectivity (Friston, 1994; Stephan and Friston, 2010)—that is, task-based synchronized activity—underlie visual-only or audiovisual speech processing, we would expect to see greater connectivity between speech-related regions during these conditions relative to auditory-only speech. In service of these questions we tested auditory-only speech perception and AV speech perception at a range of signal-to-noise ratios (SNRs) and obtained out-of-scanner measures of lipreading ability from our participants (**Figure 1**).

**Figure 1.**
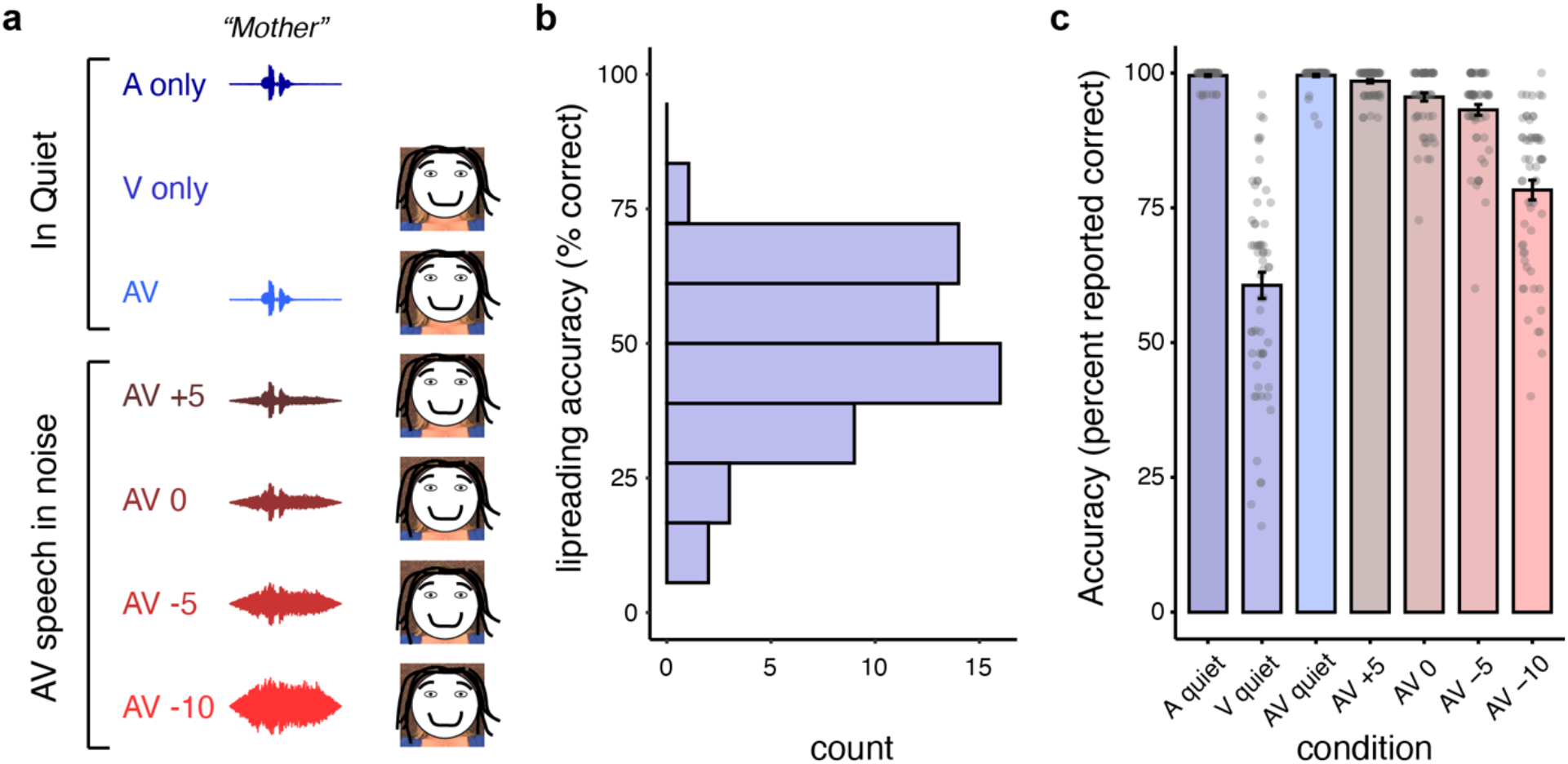
**a**. Experimental conditions with auditory-only speech, visual-only speech, and audiovisual speech. **b**. Histogram of lipreading abilities measured outside the scanner. **c**. Within-scanner behavioral performance (subjective ratings of understanding); individual participants shown in dots, mean ± SE displayed in bars.

## Method

Stimuli, behavioral data, analysis scripts, and results tables are available from https://osf.io/qxcu8/. MRI data are available on OpenNeuro (Markiewicz et al., 2021) at https://doi.org/10.18112/openneuro.ds003717.v1.0.0.

### Materials

We created seven lists of 50 words. The stimuli were recordings of a female actor speaking single words. The talker sat in front of a neutral background and spoke words along with the carrier phrase “Say the word _______” into the camera. The actor was instructed to allow her mouth to relax to a slightly open and neutral position before each target word was spoken. The edited versions of the recordings used in the current experiment did not include a carrier phrase and were each 1.5 seconds long. Recordings were made using a Canon Elura 85 digital video camera and showed the talker’s head and shoulders. Digital capture and editing were done using Adobe Premiere Elements. The original capture format for the video was uncompressed .avi; the final versions used in the study were compressed as high quality .wmv files. Audio was leveled using Adobe Audition to ensure that each word had the same root mean squared (RMS) amplitude. Conditions that included background noise used RMS-leveled six-talker babble that was mixed and included in the final version of the file.

The 350 recordings used in the study were selected from a corpus of 970 recordings of high frequency words (log HAL frequency 7.01–14.99) identified using the English Lexicon Project (Balota et al., 2007). The words that were selected for presentation in the lipreading (visual-only) or audiovisual (AV) conditions in varying signal-to-noise ratios (SNR) were selected from the larger corpus based on visual-only behavioral performance on each word from 149 participants (22–90 years old) who were tested using the entire corpus. The words selected ranged from 10%–93% correct in the lipreading-only behavioral tests. They were distributed among the six conditions that included visual information (AV in Quiet, AV +5 SNR, AV 0 SNR, AV -5 SNR, AV - 10, and visual-only) so they would, on average, be equivalent for lipreading difficulty. The words used in the auditory-only condition were selected from the remaining words.

### Participants

We collected data from 60 participants ranging in age from 18–34 years (M = 22.42, SD = 3.24, 45 female). All were right-handed native speakers of American English (no other languages other than English before age 7) who self-reported normal hearing and an absence of neurological disease. All provided informed consent under a protocol approved by the Washington University in Saint Louis Institutional Review Board.

### Procedure

Before being tested in the fMRI scanner all participants were consented, completed a safety screening, and completed an out-of-scanner lipreading assessment. The behavioral lipreading assessment consisted of 50 single word clips selected in the same way and taken from the same corpus of recorded material used in the scanner. The lipreading assessment was complete by presenting each video clip to the participant using a laptop. Participants were encouraged to verbally provide their best guess for each clip. Only verbatim responses to the stimuli were considered correct.

Participants were positioned in the scanner with insert earphones and a viewing mirror placed above the eyes to see a two-sided projection screen located at the head-side of the scanner. Those that wore glasses were provided scanner-friendly lenses that fit their prescription. Participants were also given a response box that they held in a comfortable position on their torso during testing. Each of the imaging runs presented trials with recordings of audio, visual-only, audiovisual speech stimuli, or printed text via an image projected on the screen that was visible to the participant through the viewing mirror. A camera positioned at the entrance to the scanner bore was used to monitor participant movement. A well-being check and short conversation occurred before each run and, if needed, participants were reminded to stay alert and asked to try to reduce their movement.

Six runs were completed during the session. Each run lasted approximately 5.5 minutes. The first five runs were perception runs and contained 98 trials each. The stimuli were presented in blocks of five experimental trials plus two null trials for each condition. The result was 14 blocks resulting in 70 experimental trials plus 28 null trials. All trials included 800 ms of quiet without a visual presentation before the stimuli began. During the null trials participants were presented with a fixation cross instead of the audiovisual presentation. The auditory-only condition did not include visual stimuli; instead a black screen was presented. The blocks were quasi-randomized so that two blocks from the same condition were never presented one right after the other and one null trial never occurred right after another.

To keep attention high, half of the experimental trials required a response from the participant. On response trials, a set of two dots appeared on the screen after the audiovisual/audio presentation. The right-side dot was green and the left-side dot was red. The participant was instructed to use the right-hand button on the response box to indicate “yes” if they were confident that they had been able to identify the previous word and to use the left-hand button if they felt they had not identified the previous word correctly.

After the initial five runs, a final run of 60 trials was presented in which participants saw a series of written words projected on the screen. The items were the same 50 words used for the behavioral visual-only assessment, but which did not appear in any of the other fMRI conditions. Each word stayed on the screen for 2.3 seconds, followed by two green dots that appeared for 2.3 seconds. Participants were asked to say aloud the word that was presented during the period that the dots were on the screen. Ten null trials were randomly distributed throughout the sequence. Null trials lasted 1.5 seconds and included a fixation cross on the screen. The reading task was always the final run.

### Behavioral data analysis

The out-of-scanner lipreading assessment was scored by taking the percentage of correct responses made by each participant, which we used as a covariate in the fMRI analyses, allowing us to explore patterns of brain activity that related to more successful lipreading ability. The in-scanner lipreading was scored similarly, except scores were based on participants’ own judgement of their accuracy. Because we had no way to verify lipreading accuracy in the scanner, we used these to assess qualitative differences in difficulty across condition rather than formal statistical analyses.

### MRI data acquisition and analysis

MRI images were acquired on a Siemens Prisma 3T scanner using a 32-channel head coil. Structural images were acquired using a T1-weighted MPRAGE sequence with a voxel size of .8 x .8 x .8 mm. Functional images were acquired using a multiband sequence (Feinberg et al., 2010) in axial orientation with an acceleration factor of 8 (TE = 37 ms), providing full-brain coverage with a voxel size of 2 × 2 × 2 mm. Each volume took 0.770 s to acquire. We used a sparse imaging paradigm (Edmister et al., 1999; Hall et al., 1999) with a repetition time of 2.47 s, leaving 1.7 s of silence on each trial. We presented words during this silent period, and during the repetition task, instructed participants to speak during a silent period to minimize the influence of head motion on the data.

Analysis of the MRI data was performed using Automatic Analysis version 5.4.0 (Cusack et al., 2014) (RRID:SCR_003560) that scripted a combination of SPM12 (Wellcome Trust Centre for Neuroimaging) version 7487 (RRID:SCR_007037) and FSL (FMRIB Analysis Group; Jenkinson et al., 2012) version 6.0.1 (RRID:SCR_002823). Functional images were realigned, co-registered with the structural image, and spatially normalized to MNI space (including resampling to 2 mm voxels) using unified segmentation (Ashburner and Friston, 2005) before smoothing with an 8 mm FWHM Gaussian kernel. No slice-timing correction was used. First level models contained regressors for the condition of interest (event onset times convolved with a canonical hemodynamic response function). To reduce the effects of motion on statistical results we calculated framewise displacement (FD) using the 6 realignment parameters assuming the head as a sphere with radius 50 mm (Power et al., 2012). We censored frames exceeding an FD of 0.5, which resulted in approximately 8% data loss across all participants. Frames with FD values exceeding this threshold were modeled out by adding in one additional column to the design matrix for each high-motion scan (cf. Lemieux et al., 2007).

Psycho-physiological interaction (PPI) analyses are designed to estimate the effective connectivity between brain regions (Friston et al., 1997); that is, the degree to which task demands alter the functional connectivity (i.e., statistical dependence of time series) between a seed region and every other voxel in the brain. PPI analyses thus require identifying a seed region from which to extract a time course, and two (or more) tasks between which to compare connectivity with the seed region. For auditory and visual cortex ROIs (see below for definition), we extracted the time course of the seed region using SPM’s VOI functionality, summarizing the time course as the first eigenvariate of the ROI after adjusting for effects of interest.

Contrast images from single subject analyses were analyzed at the second level using permutation testing (FSL *randomise;* 5000 permutations) with a cluster-forming threshold of p < .001 (uncorrected) and results corrected for multiple comparisons based on cluster extent (p < .05). Anatomical localization was performed using converging evidence from author experience (Devlin and Poldrack, 2007) viewing statistical maps overlaid in MRIcroGL (Rorden and Brett, 2000), supplemented by atlas labels (Tzourio-Mazoyer et al., 2002).

### Regions of interest

We defined regions of interest (ROIs) for the left posterior temporal sulcus (pSTS), left primary auditory cortex (A1), and left primary visual cortex (V1). For the pSTS, the ROI was defined as a 10 mm radius sphere centered at MNI coordinates (x=-54, y=-42, z=4) previously reported to be activated during audiovisual speech processing (Venezia et al., 2017). The ROIs for AI and V1 were defined using the Anatomy Toolbox (Eickhoff et al., 2005) (RRID:SCR_013273) as the combination of Areas TE1.0, TE 1.1, and TE 1.2 in the left hemisphere (Morosan et al., 2001) and the left half of area hOC1, respectively. For the non-PPI ROI analysis, data were extracted by taking the mean of all voxels in each ROI.

## Results

Unthresholded statistical maps are available from NeuroVault (Gorgolewski et al., 2015) at https://neurovault.org/collections/10922/.

We first examined whole brain univariate effects by condition, shown in **Figure 2**. We observed temporal lobe activity in all conditions, including visual-only, and visual cortex activity in all conditions except auditory only.

**Figure 2.**
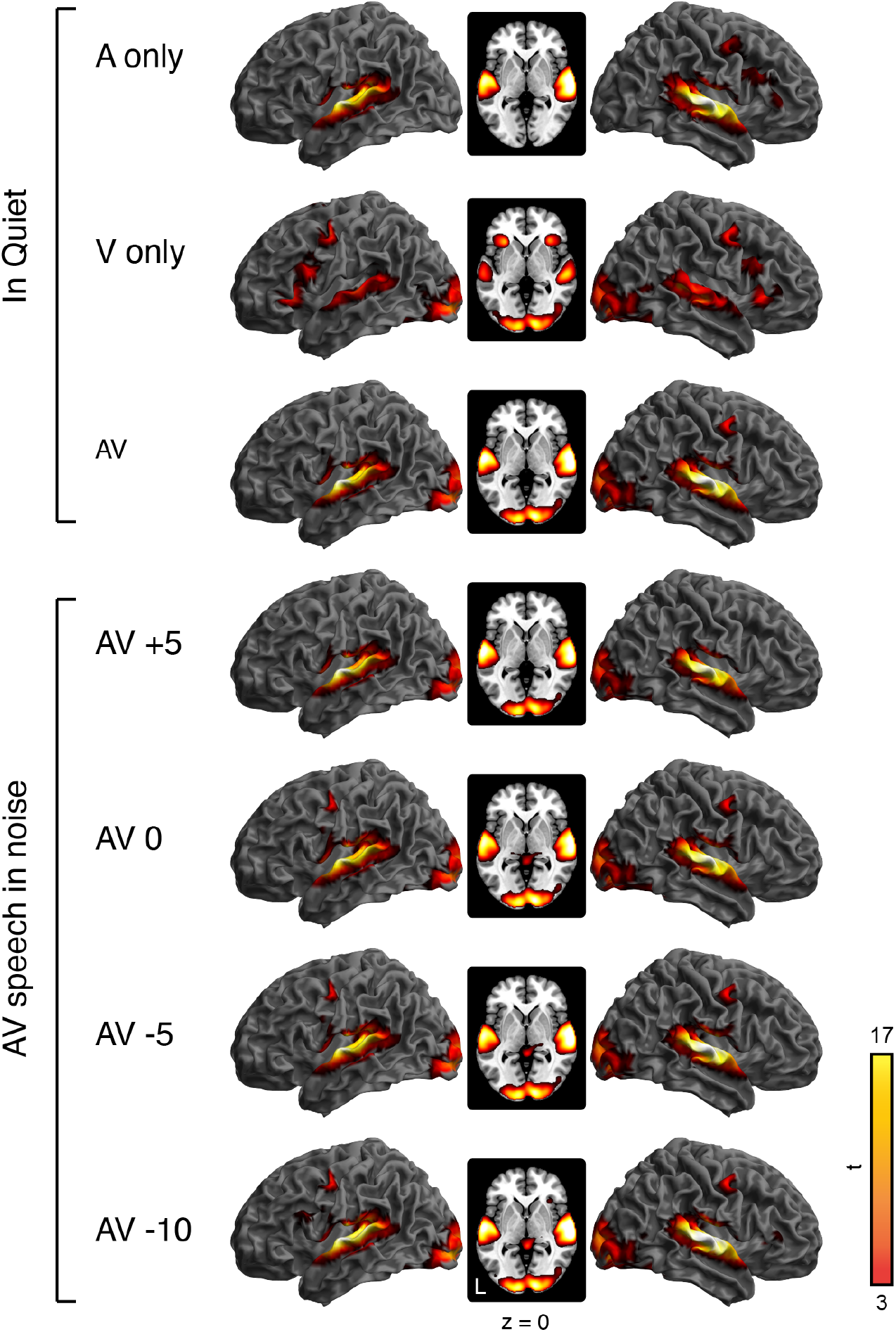
Univariate results for spoken word perception in all experimental conditions.

We next related the activity during visual-only speech with the out-of-scanner lipreading score (**Figure 1b**). Across participants, lipreading accuracy ranged from 4– 74% (mean = 47.75, SD = 15.49), and correlated with in-scanner ratings (Spearman rho = 0.38). We included out-of-scanner lipreading as a covariate to see whether individual differences in out-of-scanner scores related to visual-only activity; we did not find any significant relationship (positive or negative).

Following univariate analyses, we examined effective connectivity using psychophysiological interaction (PPI) models. We started by using a seed region in left visual cortex. As seen in **Figure 3**, compared to auditory-only speech, visual-only and all audiovisual conditions showed increased connectivity with the visual cortex seed, notably including bilateral superior temporal gyrus and auditory cortex. The same was true with an auditory cortex seed, shown in **Figure 4**. Here, compared to the visual-only condition, we see increased connectivity with visual cortex in all conditions except the auditory-only condition.

**Figure 3.**
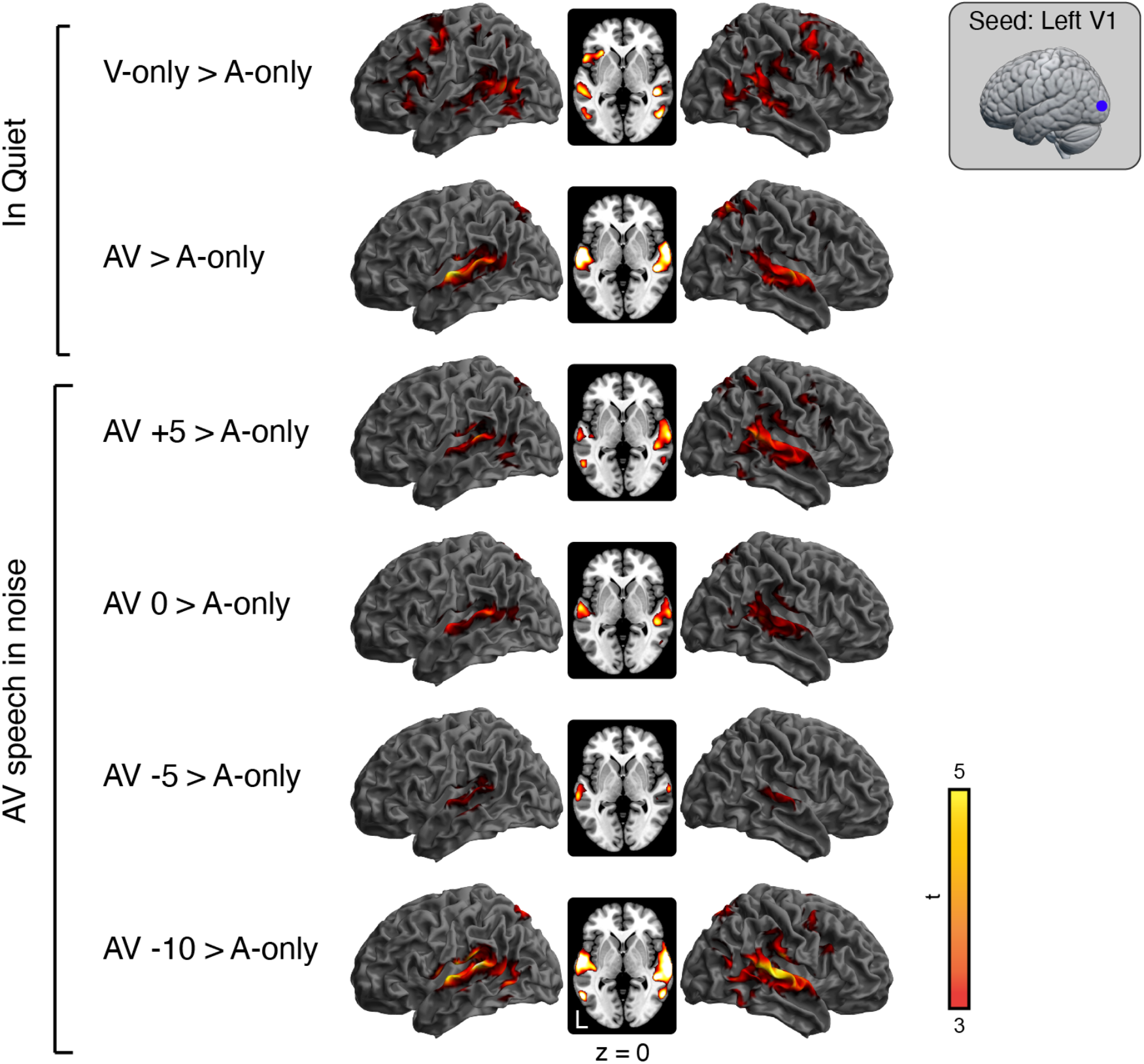
Psychophysiological interaction analysis for experimental conditions, using a seed from left visual cortex. Warm-colored voxels showed significantly more connectivity with visual cortex in an experimental condition than in the auditory-only condition.

**Figure 4.**
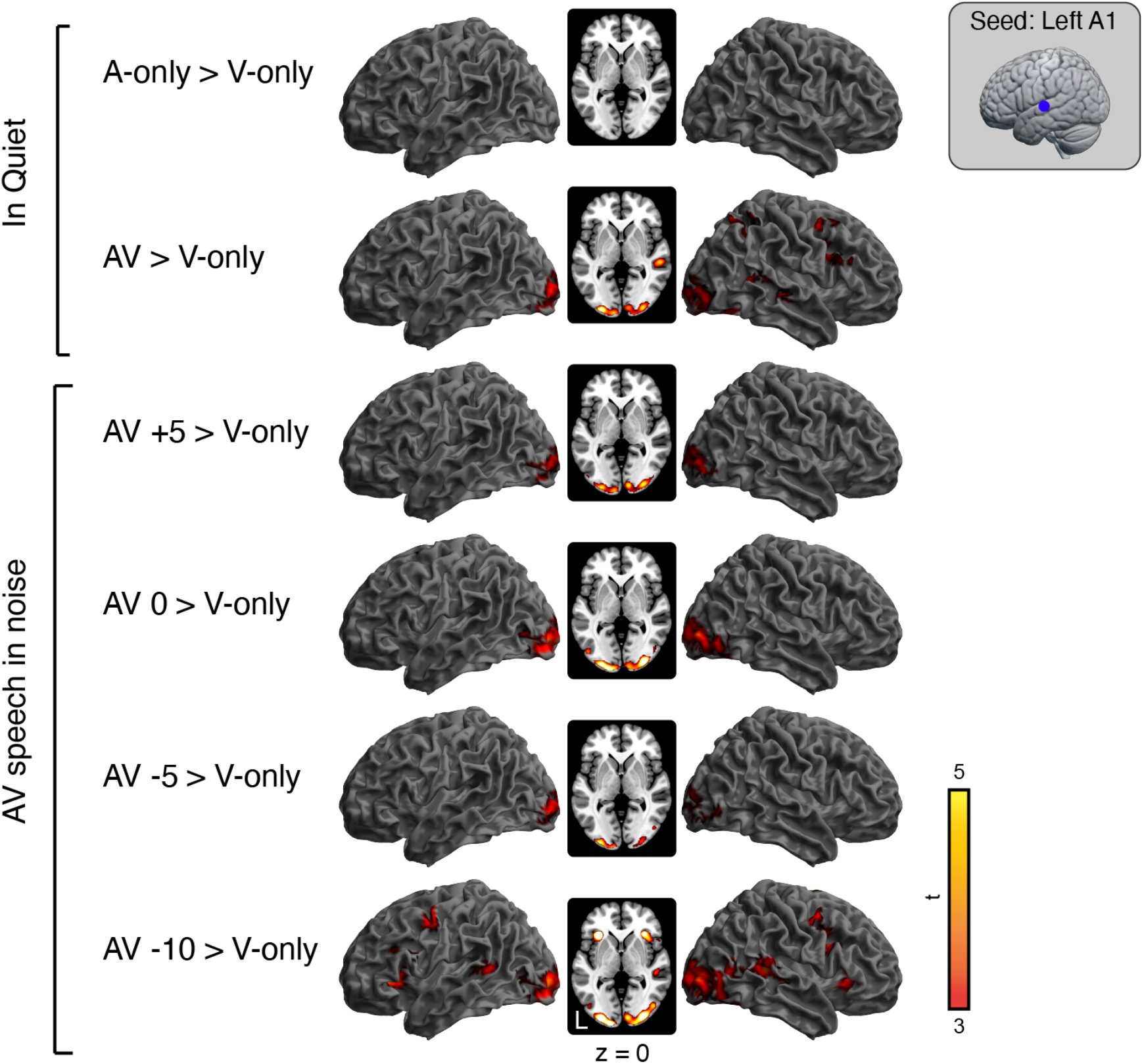
Psychophysiological interaction analysis for experimental conditions, using a seed from left auditory cortex. Warm-colored showed significantly more connectivity with auditory cortex in an experimental condition than in the visual-only condition.

Finally, to complement the above whole-brain analyses, we conducted an ROI analyses focusing on pSTS, shown in **Figure 5**. For the whole-brain univariate and PPI analyses described above, we extracted values from left pSTS and used one-sample t-tests to see whether activity was significantly different from 0. Significance (p < .05, Bonferroni corrected for 19 tests giving p < .00263) is indicated above each condition.

**Figure 5.**
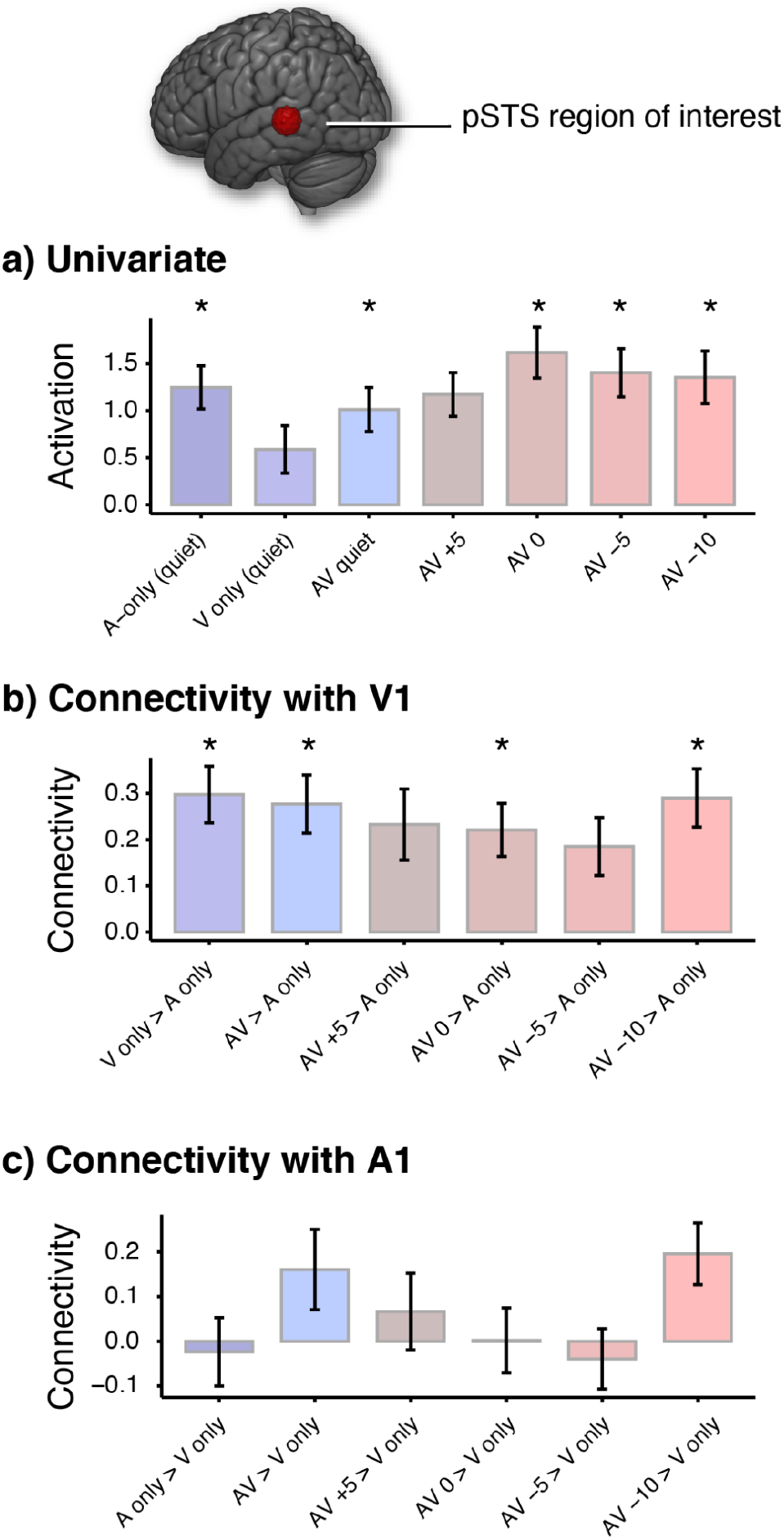
Region-of-interest analyses highlighting the role of the left pSTS in speech processing. **a**. pSTS activity for univariate analyses (cf. Figure 2). **b**. PPI-based effective connectivity with V1 (cf. Figure 3). **c**. PPI-based effective connectivity with A1 (cf. Figure 4). Significant differences from 0, corrected for multiple comparisons, are indicated with an asterisk.

## Discussion

We studied brain activity associated with visual-only and audiovisual speech perception. We found that connectivity between auditory, visual, and premotor cortex was enhanced during audiovisual speech processing relative to unimodal processing, and during visual-only speech processing relative to auditory-only speech processing. These findings are broadly consistent with a role for synchronized interregional neural activity supporting visual and audiovisual speech perception.

### Dedicated regions for multisensory speech processing

Although understanding audiovisual speech requires combining information from multiple modalities, the way this happens is unclear. One possibility is that heteromodal brain regions such as the pSTS act to integrate inputs from unisensory cortices. In addition to combining inputs to form a unitary percept, regions such as pSTS may also give more weight to more informative modalities (for example, to the visual signal when the auditory signal is noisy) (Nath and Beauchamp, 2011).

Activity in pSTS for visual-only or AV speech was suggested by both our whole-brain and ROI-based analyses. In particular, we observed pSTS activity for AV speech in which the auditory and visual aspects were consistently congruent, consistent with a role for pSTS in integrating or combining auditory and visual information. Of course, pSTS activity is not always observed for AV speech (Erickson et al., 2014). One potential explanation for the variability in pSTS activation across studies is nature of the speech materials. Several previous studies identifying pSTS involvement in multisensory speech perception have used incongruent stimuli (i.e., a McGurk task) (McGurk and MacDonald, 1976), which differs substantially from most of our everyday speech perception experience (Van Engen et al., 2019). Thus, the conditions under which pSTS is recruited to support visual or AV speech perception remains an open question.

In our univariate results, we observed activity in premotor cortex for both visual-only speech in quiet, and AV speech at more challenging signal-to-noise ratios. These findings are consistent with a flexible role for premotor cortex in speech perception, at least under some circumstances, as reported in other studies of visual and audiovisual speech perception (Venezia et al., 2017). Although our current data do not support specific conclusions, the dependence of premotor activity on task demands may explain some of the inconsistencies underlying the debates about the role of premotor cortex that permeate the speech perception literature.

### Effective connectivity and multisensory speech processing

A different perspective comes from a focus on multisensory effects in auditory and visual cortex (Peelle and Sommers, 2015). Much of the support for this “early integration” view comes from electrophysiology studies showing multimodal effects in primary sensory regions (e.g., Schroeder and Foxe, 2005). For example, Lakatos and colleagues (2007) found that somatosensory input reset the phase of ongoing neural oscillations in auditory cortex, which was hypothesized to increase sensitivity to auditory stimuli. In at least one human MEG study, audiovisual effects appear sooner in auditory cortex than in pSTS (Möttönen et al., 2004), and visual speech may speed processing in auditory cortex (van Wassenhove et al., 2005). These findings suggest that multisensory effects are present in primary sensory regions, and that auditory and visual information do not require a separate brain region in which to “integrate”.

In the current data, we observed stronger connectivity between auditory and visual cortex for visual-only and audiovisual speech conditions than for unimodal auditory-only speech; and stronger connectivity in audiovisual speech conditions than in unimodal visual-only speech. That is, using a visual cortex seed we found increases in effective connectivity with auditory cortex, and when using an auditory cortex seed we found increases in effective connectivity with visual cortex. These complementary findings indicate that functionally coordinated activity between primary sensory regions that is increased during audiovisual speech perception.

Beyond primary sensory cortices, we also observed effective connectivity changes to premotor cortex for both visual-only speech and several audiovisual conditions. The functional synchronization between visual cortex, auditory cortex, and premotor cortex is consistent with a distributed network that orchestrates activity in response to visual-only and audiovisual speech.

Finally, our ROI analysis showed increased effective connectivity between pSTS and V1, but not A1, under most experimental conditions (**Figure 5**). These effective connectivity changes with V1 are consistent with a role for pSTS in audiovisual speech processing. However, they are also not easily reconcilable with studies reporting connectivity differences between pSTS and both A1 and V1 (Nath and Beauchamp, 2011). Although no doubt the location and size of any pSTS ROI chosen is important, we used the same ROI for the PPI analyses with both the A1 seed and V1 seed, and so ROI definition alone does not seem to explain the qualitative difference between the two.

It may be worth considering whether the pSTS plays different role in relation to A1 and V1. Just because pSTS responds to both auditory and visual information does not necessarily mean it treats them equally, or integrates them in a modality-agnostic manner. Indeed, given that “unisensory” cortices show multisensory effects and anatomical connections (Cappe & Barone, 2005), heteromodal or multisensory regions can also exhibit modality preferences (Noyce et al., 2017). In many audiovisual tasks, auditory information appears to be preferentially processed (Grondin and McAuley, 2009; Grondin and Rousseau, 1991; Grahn et al. 2011; Recanzone, 2003). Thus, pSTS may be particularly important in integrating visual information into an existing auditory-dominated percept. Relatedly, it could also be that multimodal information is inextricably bound at early stages of perception (Rosenblum, 2008), a process which may rely on pSTS.

The emerging picture is one in which coordination of large-scale brain networks—that is, effective connectivity reflecting time-locked functional processing—is associated with visual-only and audiovisual speech processing. What might be the function of such distributed, coordinated activity? Visual and audiovisual speech appear to rely on multisensory representations. For audiovisual speech, it may seem obvious that successful perception requires combining auditory and visual information. However, visual-only speech has been consistently associated with activity in auditory cortex (Calvert et al., 1997; Okada et al., 2013). These activations may correspond to visual-auditory associations, and auditory-motor associations, learned from audiovisual speech that are automatically reactivated, even when the auditory input is absent.

Interestingly, our out-of-scanner lipreading scores did not correlate with any of the whole brain results. It should be noted, however, that our sample size, while large for fMRI studies of audiovisual speech processing, may still be too small to reliably detect individual differences in brain activity patterns (Yarkoni and Braver, 2010). Moreover, there may be multiple ways that brains can support better lipreading, and such heterogeneity in brain patterns would not be evident in our current analyses. Future studies with larger sample sizes may be needed to quantitatively assess the degree to which users’ activity might fall into neural strategies, and the degree to which these are related to lipreading performance.

It is worth highlighting an intriguing aspect of our data, which is that auditory cortex is always engaged, even in visual-only conditions, whereas the reverse is not true for visual cortex (which is only engaged when visual information is present) (**Figure 2**). This observation may relate to deeper theoretical issues regarding the fundamental modality of speech representation. That is, if auditory representations have primacy (at least, for hearing people), we might expect these representations to be activated regardless of the input modality (i.e., for both auditory and visual speech). In fact, this is exactly what we have observed. Although these findings do not directly speak to the level of detail contained in visual cortex speech representations (Bernstein and Liebenthal, 2014), they are consistent with asymmetric auditory and visual speech representations.

### Different perspectives on multisensory integration during speech perception

An enduring challenge for understanding multisensory speech perception can be found in differing uses of the word “integration”. During audiovisual speech perception, listeners use both auditory and visual information, and so from one perspective both kinds of information are necessarily “integrated” into a listener’s (unified) perceptual experience. However, such use of both auditory and visual information does not necessitate a separable cognitive stage for integration (Tye-Murray et al., 2016; Sommers, 2021), nor does it necessitate a region of the brain devoted to integration. The interregional coordination we observed here may accomplish the task of integration in that both auditory and visual modalities are shaping perception. In this framework, there is no need to first translate visual and auditory speech information into some kind of common code (see also Altieri et al., 2011).

With any study it is important to consider how the specific stimuli used influenced the results. Here, we examined processing for single words. Visual speech can inform perception in multiple dimensions (Peelle and Sommers, 2015), including by providing clues to the speech envelope (Chandrasekaran et al., 2009). These clues may be more influential in connected speech (e.g., sentences) than in single words, as other neural processes may come into play with connected speech.

## Conclusion

Our findings demonstrate the scaffolding of connectivity between auditory, visual, and premotor cortices that supports visual-only and audiovisual speech perception. These findings suggest that the binding of multisensory information need not be restricted to heteromodal brain regions (e.g., pSTS), but may also emerge from coordinated unimodal activity throughout the brain.

## Acknowledgments

This work was supported by grants R56 AG018029 and R01 DC016594 from the US National Institutes of Health. The multiband echo planar imaging sequence was provided by the University of Minnesota Center for Magnetic Resonance Research.

## Notes

**Conflicts of interest:** The authors declare no competing financial conflicts of interest.

https://osf.io/qxcu8/

https://doi.org/10.18112/openneuro.ds003717.v1.0.0

### Competing Interest Statement

The authors have declared no competing interest.

